# Integration of genomic variation and phenotypic data using HmtPhenome

**DOI:** 10.1101/660282

**Authors:** R. Preste, M. Attimonelli

**Affiliations:** Department of Biosciences, Biotechnologies and Biopharmaceutics, University of Bari, Bari 70126, Italy

## Abstract

A full understanding of relationships between variants, genes, phenotypes and diseases is often overlooked when investigating mitochondrial functionality in both healthy and pathological situations. Gaining a comprehensive overview of this network can indeed offer interesting insights, and guide researchers and clinicians towards a full-spectrum knowledge of the mitochondrial system.

Given the current lack of tools addressing this need, we have developed HmtPhenome (https://www.hmtphenome.uniba.it), a new web resource that aims at providing a visual network of connections among variants, genes, phenotypes and diseases having any level of involvement in the mitochondrial functionality. Data are collected from several third party resources and aggregated on the fly, allowing users to clearly identify interesting relations between the involved entities. Tabular data with additional hyperlinks are also included in the output returned by HmtPhenome, so that users can extend their analysis with further information from external resources.

## Introduction

The relationship between mitochondria and disease has gained a lot of interest by clinicians and researchers during recent years, due to the involvement of the mitochondrion in several biological processes, the most important being aerobic ATP production. Defective mitochondria can be, directly or indirectly, associated to the onset and progression of neurodegenerative diseases^1–3^, diabetes^4^, cancer^5, 6^, metabolic syndromes^7^ and other pathologies, with a broad spectrum of phenotypic traits and outcomes.

Moreover, human cells present a high number of mitochondria, each one of which can contain thousands of copies of mtDNA with a variable ratio of wild-type/mutated genomes, a condition defined as mitochondrial heteroplasmy. Aberrant mitochondrial phenotypes can only arise when the number of mutated genomes is sufficient to take over the wild type ones and exert their detrimental effects. This can explain the pathologic heterogeneity usually observed in mitochondrial diseases: the same disease can show slightly different phenotypes among different individuals, based on the heteroplasmic fraction of the given causative mutation(s). Recent clinical literature has shown several examples of pathologies characterised by different levels of severity or phenotypic effects due to variable mitochondrial heteroplasmic fraction^8–10^.

Furthermore, mitochondrial dysfunction can also occur when mutations are located in nuclear-encoded genes whose functionality affects that of mitochondria. More than 1000 genes exist that are coded by the nuclear genome but are involved in mitochondrial physiology, and most of them are implicated in energy production processes at various levels. A comprehensive list of such genes is maintained by Mitocarta^11^, which provides an index of 1158 genes involved in mitochondrial activities.

Many online resources offer data about mitochondrial variations and their involvement in pathological conditions, such as Mitomap^12^, LOVD^13^, HmtDB^14^, HmtVar^15^, Mitobreak^16^, MitImpact^17^. However, while these softwares extensively analyse mitochondrial mutations at different levels, from large genomic rearrangements to SNPs, they usually only report diseases related to a specific variation, lacking to properly highlight potential relationships between variations and their phenotypic effects. These information can indeed be of much help for researchers and clinicians to obtain a much broader view of the studied subject.

An additional source of difficulty that is frequently encountered when investigating phenotypes and diseases is represented by the different ways in which the same condition can be potentially referred to. Phenotype and disease names can differ over distinct resources, and the actual cut-off line between these two entities is often blurred, confusing phenotypic effects with disease causing them, and vice-versa. This issue was effectively addressed by ontology services, which offer standardised vocabularies that unambiguously identify entities and relationships among them^18^. Terms are arranged in a hierarchical manner, with terms referring to much broader concepts located as top nodes, and more specific elements can be found traversing this tree-like structure until reaching the final leaves; each node usually contains other additional information about the described element, with links pointing to further external resources. The Human Phenotype Ontology (HPO)^19^ and the Disease Ontology (DO)^20^ are probably the most commonly used examples of such schema as related to phenotypes and diseases, respectively, but many other similar services exist, like the Mammalian Phenotype Ontology (MPO)^21^, PhenomeNET^22^, Medical Subject Headings (MeSH)^23^ and the Unified Medical Language System (UMLS)^24^.

In order to better understand the pathological mechanisms of mitochondrial syndromes and diseases, it is then fundamental to build a comprehensive network of diseases and related phenotypes directly or indirectly associated to mitochondrial mutations; along with them, nuclear and mitochondrial genes with some level of involvement in mitochondrial functionality should be considered as well.

For this reason we developed HmtPhenome (https://www.hmtphenome.uniba.it), a system that integrates information about variants, genes, phenotypes and diseases associated to mitochondrial functions, and allows to perform queries that can start from any one of these entry points, retrieving data related to the other three entities and building a full-fledged information network.

## Materials and methods

The aim of HmtPhenome is to provide information about four different biological entities and the relationship among them, namely variants, genes, phenotypes and diseases with any involvement in mitochondrial functionality. In order to do this, HmtPhenome retrieves the relevant data from several online resources, interconnects these information when possible and returns a network that highlights relations of interest for researchers and clinicians.

Due to the high number of resources involved and the great amount of data considered in each query, a classic data retrieval system using a local database would have not been suitable for these purposes; instead, all the needed information is collected from external resources on the fly after a user query, and only a small amount of data is actually stored in a local database. The local database is mostly used by the web interface to populate query menus and provide autocompletion suggestions based on user input, thanks to the Awesomplete Javascript framework^25^. This set of locally-stored information contains:

- the list of 1158 mitochondrial- and nuclear-encoded genes involved in mitochondrial functionality, collected from Mitocarta and used in the Gene query section; this list is further extended with the 22 mitochondrial tRNA genes, totalling 1180 genes;
- a list of diseases collected from OMIM^26^ and Orphanet^27^, used to provide suggestions in the Disease query section; this basic list is further enriched using data coming from DisGeNET^28^, to integrate additional information about variants and genes associated with each disease;
- a list of phenotypes, collected from HPO and used to provide suggestions in the Phenotype query section; additional data are also collected from HPO with details about the association of these phenotypes with specific diseases and genes;
- a list of mitochondrial variants derived from HmtVar, which are used to shorten waiting times when querying the system for variations occurring on the mitochondrial genome.

All the other information related to variants, genes, phenotypes and diseases and their mutual relationships are gathered from third-party resources, exploiting their APIs, such as HPO, the Experimental Factor Ontology (EFO)^29^, Ensembl REST services^30^ and BioMart^31^.

Having to deal with a large amount of data flowing in from several external services, there is the risk that the system may face long-running tasks when waiting for their response, and thus users could experience delays while using HmtPhenome. This possibility was taken into account and addressed using complementary strategies: first of all, HmtPhenome was built using the Quart Python framework^32^, which is specifically suited to handle all the requests to external services separately and in parallel, instead of queueing them one after the other, reducing waiting times sensibly. Furthermore, HmtPhenome uses a caching system that avoids repeated requests to external resources in case of the same query performed more than once in a short time span: in this case, results are retrieved and returned from the local cache memory instead, further shortening waiting times. In addition, in case third-party resources take too long or fail to provide a valid response, a fallback set of basic information retrieved from the local database is returned, so that a minimum amount of data is always available in the query results.

The final data, collected from both the local database and external resources, are aggregated to create a dictionary-like structure, where the keys are represented by the above-mentioned four different biological entities, i.e. disease, genes, variants and phenotypes, and the value associated to each key is a list of all the available information in that context. The key referring to the query starting point will obviously contain a single element, the query subject, with the only exception being when the query starts from a variant position, in which case multiple variants can be found on the same genomic position. This dictionary structure is rendered as both a visual network and a tabular form, thanks to the vis.js^33^ and DataTables^34^ Javascript frameworks, respectively. In addition, a JSON-formatted file and a MS Excel tabular file containing the same data are also provided and available for download, for users that may want to perform further downstream analyses.

## Results and discussion

HmtPhenome offers a powerful Query page, where users can select which biological entity they want to focus on, and retrieve all the available information about it.

Queries on HmtPhenome are organised on different levels, and can start from one of these entry points: variant position;

- variant position and optionally alternate allele;
- gene;
- phenotype;
- disease.

In general, when a query is launched on HmtPhenome, a basic set of information about the chosen biological entity is first retrieved from the local database, depending on the specific entry point; then, API requests are performed to the relevant external services, in order to collect as many additional data as possible.

All these information are finally integrated to build the final table listing all the different variants, genes, phenotypes and diseases found, with links pointing to external resources with more information about each of them; the same data can also be visualised through a network shown on a separate page, where every different piece of information available is connected to another according to their biological relationships; variants, genes, phenotypes and diseases are shown in different colours to facilitate the visual recognition of interesting relations and patterns in this network. JSON-formatted and MS Excel tabular files with the same data are also available for further manipulation.

For queries starting from a variant, users will first have to select a specific chromosome to restrict the search; after that, users can type a specific position to investigate on that chromosome and an alternate allele to further limit their search. After the query is launched, one or more variants (depending whether the query involved only the variant position or also a specific allele) are found and connected to the gene they belong to (if applicable), to the diseases they are involved in and to the phenotypes they cause or the phenotypes that may cause the disease, when these information are available (Fig. 1).

**Fig. 1.**
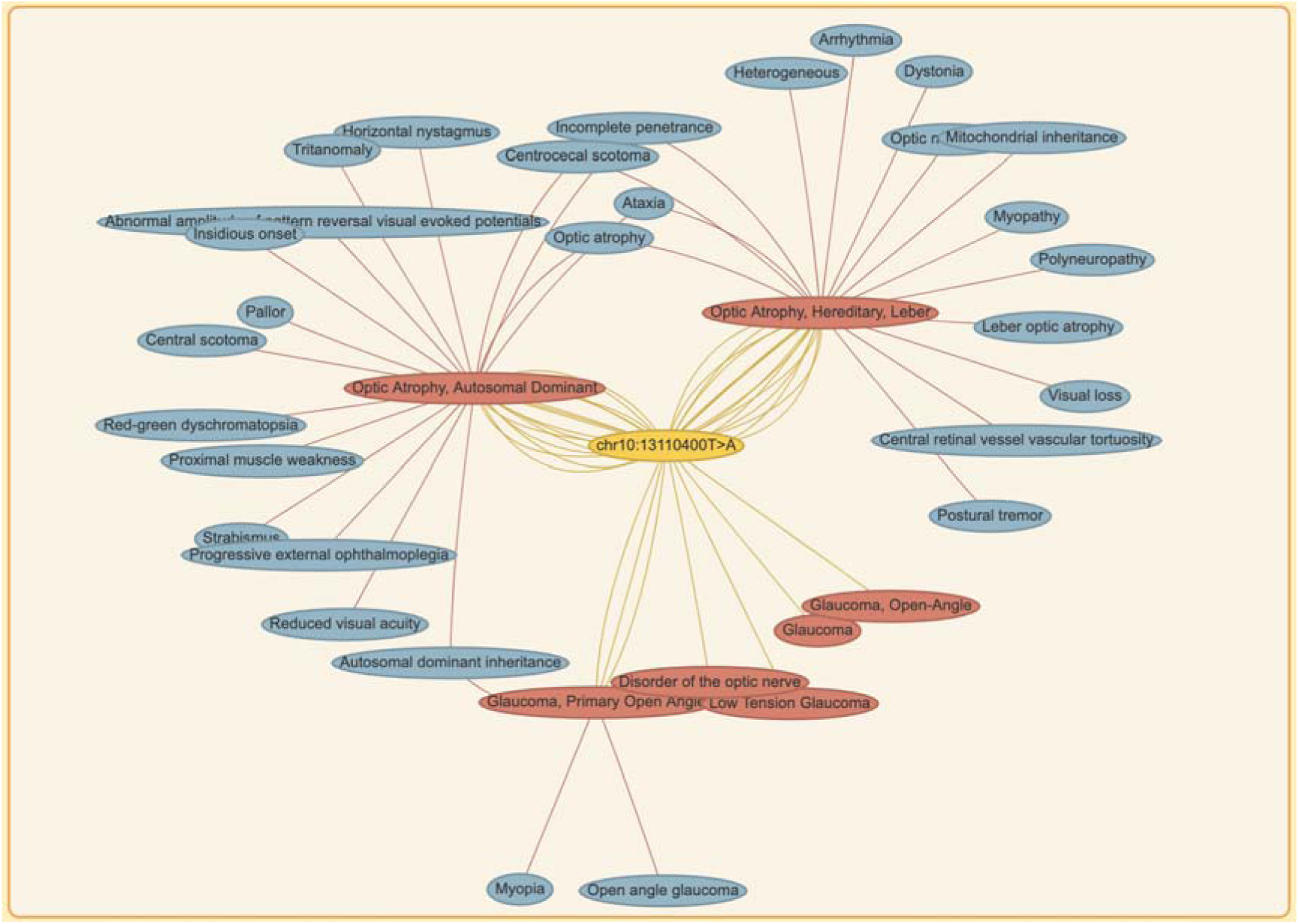
Network returned after a query by variant position (chr10:13110400T>A). Yellow nodes: variants, green nodes: genes (not shown here), red nodes: diseases, blue nodes: phenotypes.

If users want to investigate a specific gene, they can insert the desired Ensembl Gene ID or start typing the common gene name in the related text search area, and a few suggestions from the list of Mitocarta genes will be shown to facilitate the search. Query results report variants associated with the chosen gene, diseases and phenotypes related to these variants as well as those related to the queried gene, when further information about involved variants is not available (Fig. 2).

**Fig. 2.**
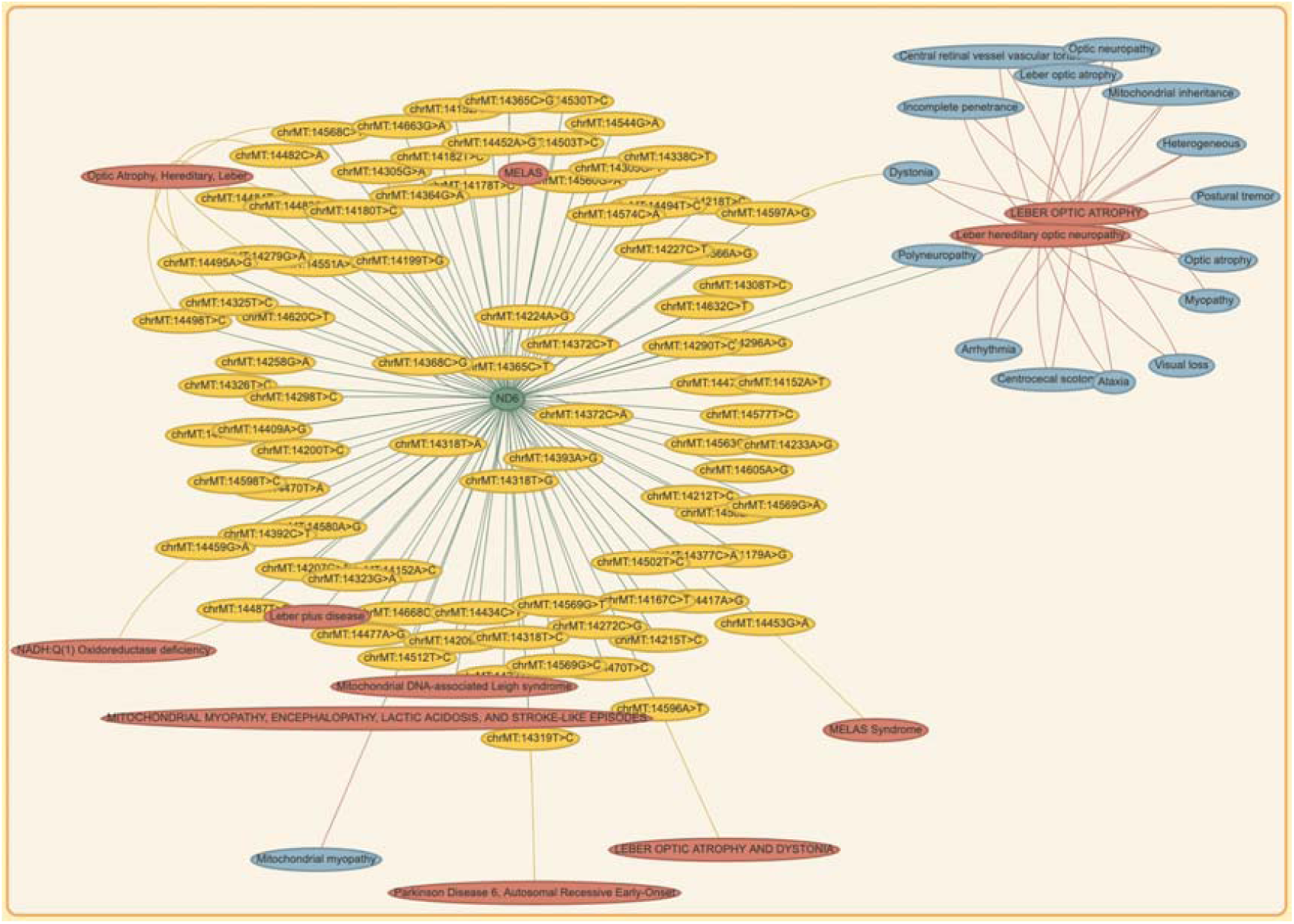
Network returned after a query by gene name (ND6, Ensembl Gene ID ENSG00000198695). Yellow nodes: variants, green nodes: genes, red nodes: diseases, blue nodes: phenotypes.

Queries involving diseases also present an open text search, which can accept either Orphanet or OMIM identifiers. As an alternative, users can also start typing the desired common disease name, and a set of related suggestions will show up to guide the user; when a suggested element is selected, the associated identifier will be typed in automatically. Disease queries return information about genes and variants known to be involved in the given disease onset or progression, as well as phenotypes caused by or somewhat associated to it (Fig. 3). The query disease name is also converted to its proper ontology term, according to the UMLS service.

**Fig. 3.**
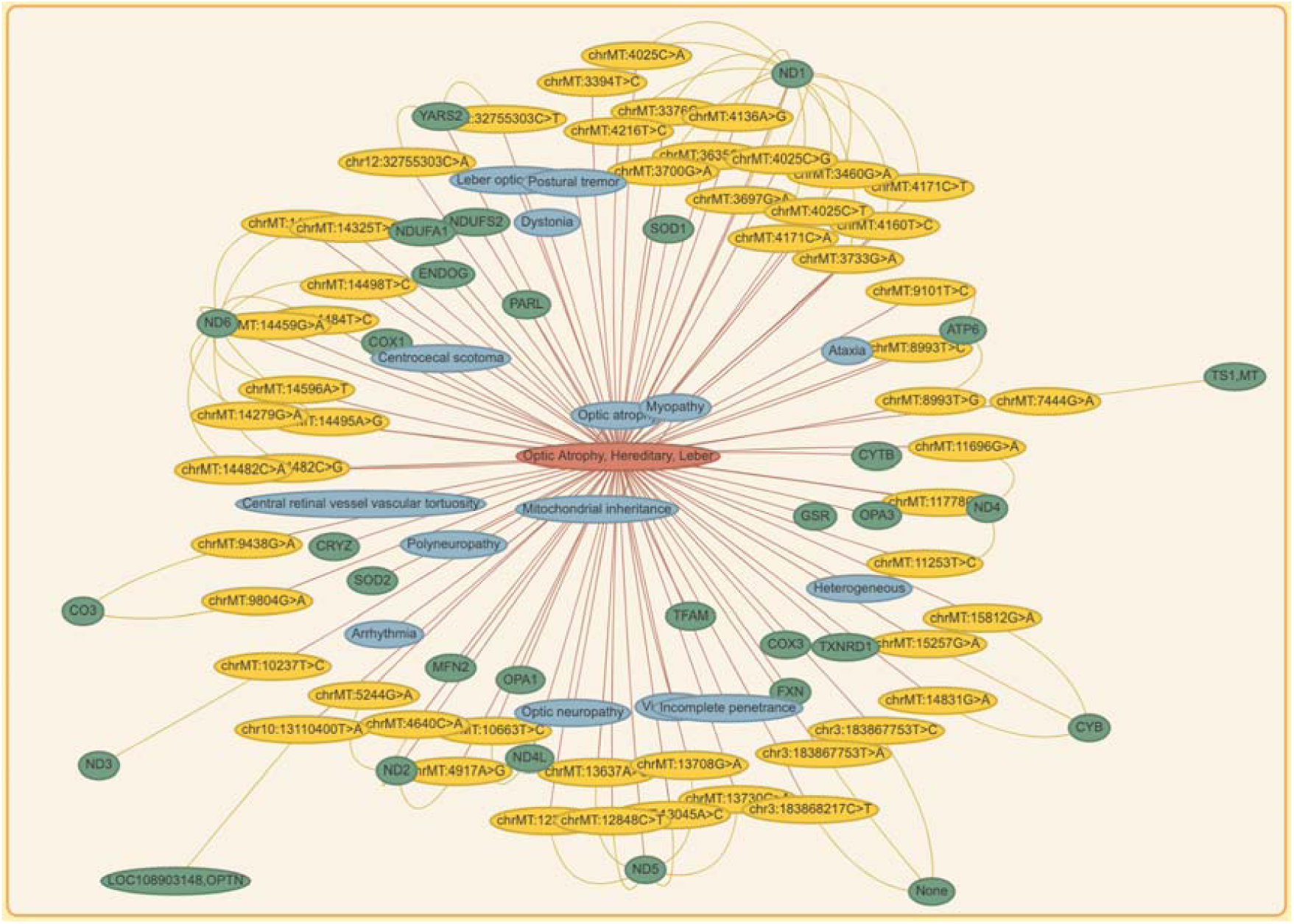
Network returned after a query by disease name (Leber Optic Atrophy, OMIM:535000). Yellow nodes: variants, green nodes: genes, red nodes: diseases, blue nodes: phenotypes.

The query field for phenotypes also features an open text search with autocompletion, but in this case it accepts HPO identifiers related to phenotype names. Query results contain diseases characterised by the given phenotype, together with genes and variants that are related to it (Fig. 4).

**Fig. 4.**
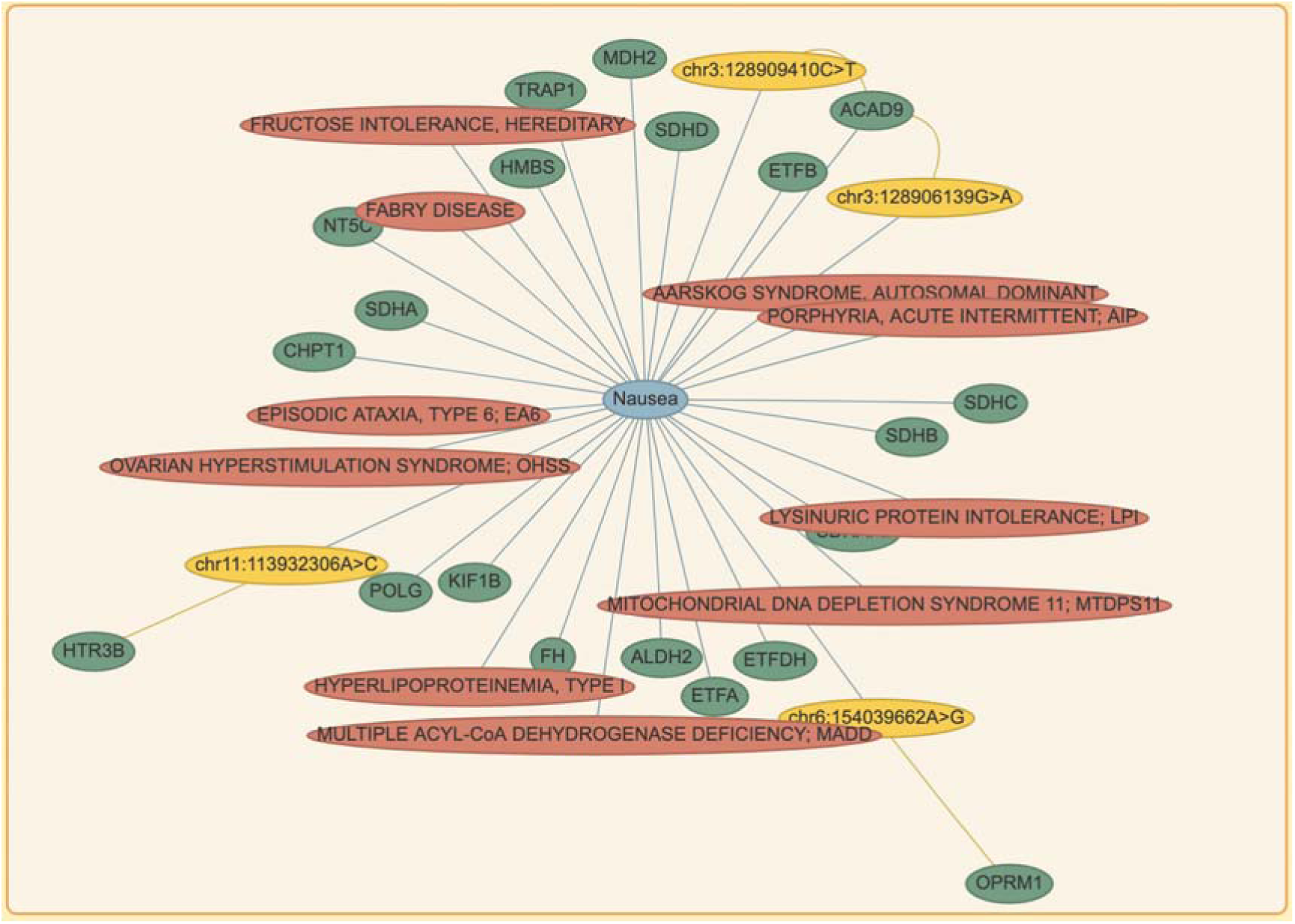
Network returned after a query by phenotype name (Nausea, HP:0002018). Yellow nodes: variants, green nodes: genes, red nodes: diseases, blue nodes: phenotypes.

## Conclusions

HmtPhenome (https://www.hmtphenome.uniba.it) offers a comprehensive view of relationships between variants, genes, phenotypes and diseases with a particular involvement in mitochondrial functionality. These data can be extremely useful for researchers and clinicians looking to integrate information coming from different resources and focusing on separate aspects of physiological and pathological mitochondria.

Given the high number of resources involved and the impressive amount of data generated and manipulated by HmtPhenome, a great effort was made to ensure this system would work seamlessly and efficiently, through the usage of state-of-the-art programming frameworks and a set of default fallback data. The most useful online resources with data about variants, gene, diseases and phenotypes are scanned to gather information about these biological entities, and these information are then thoroughly aggregated and returned to the user both in graphical and textual form. This allows to identify and further investigate existent biological relationships at a glance, as well as finding potential new players involved in mitochondrial functionality to some extent.

Although the current implementation of HmtPhenome queries a limited number of third-party resources, the returned set of information is extensive and serves efficiently its purpose of providing a wide overview of the relationships existing among variants, genes, diseases and phenotypes, particularly as regarding mitochondrial functionality. More external resources will be integrated into this system with the upcoming updates, with particular attention to mitochondria-focused ones, in order to increase the amount of data processed and augment the added value of the information integration performed by HmtPhenome.

